# Functional repurposing of regulatory element activity during mammalian evolution

**DOI:** 10.1101/251371

**Authors:** Francesco N. Carelli, Angélica Liechti, Jean Halbert, Maria Warnefors, Henrik Kaessmann

## Abstract

The spatiotemporal control of gene expression exerted by promoters and enhancers is central for organismal development, physiology and behaviour. These two types of regulatory elements have long been distinguished from each other based on their function, but recent work highlighted common architectural and functional features. It also suggested that inheritable alterations in the epigenetic and sequence context of regulatory elements might underlie evolutionary changes of their principal activity, which could result in changes in the transcriptional profile of genes under their control or even facilitate the birth of new genes. Here, based on integrated cross-mammalian analyses of DNase hypersensitivity, chromatin modification and transcriptional data, we provide support for this hypothesis by detecting 449 regulatory elements with signatures of activity turnover in sister species from the primate and rodent lineages (termed “P/E” elements). Through the comparison with outgroup species, we defined the directionality of turnover events, which revealed that most instances represent transformations of ancestral enhancers into promoters, leading to the emergence of species-specific transcribed loci or 5’ exons. Notably, P/E elements have distinct GC sequence compositions and stabilizing 5’ splicing (U1) regulatory motif patterns, which may predispose them to functional repurposing during evolution. Moreover, we trace changes in the U1 and polyadenylation signal densities and distributions that accompanied and likely drove the evolutionary activity switches. Overall, our work suggests rather widespread evolutionary remodelling of regulatory element functions. Functional repurposing thus represents a notable mechanism that likely facilitated regulatory innovation and the origination of new genes and exons during mammalian evolution.

## INTRODUCTION

Gene transcription in mammals is controlled by the complex interactions between different classes of proximal and distal regulatory elements. Promoters – the proximal cis-regulatory regions associated to the transcription start site (TSS) of a gene – mediate the recruitment of the RNA polymerase II (Pol II) through their recognition by general transcription factors^1^. The correct spatio-temporal activation of gene expression is further defined by transcription factors bound to other classes of regulatory loci, including TSS-distal enhancers^2,3^. Analyses of individual regulatory elements and genome-wide surveys revealed sequence and structural features characterizing promoters and enhancers in a number of species. Most vertebrate promoters are CpG-rich^1^, while most enhancers are CpG-poor^4^, a difference that is also reflected in the respective regulatory motif compositions^5,6^. Furthermore, while both of these elements are characterized by accessible chromatin, as revealed by analyses of genome-wide DNase hypersensitivity sequencing (DNase-seq) data^7^, enhancers and promoters can be distinguished based on different chromatin modification profiles. Promoters are associated with higher levels of trimethylation of lysine 4 at histone 3 (H3K4me3) compared to monomethylation of the same residue (H3K4me1), whereas the opposite pattern is found for enhancers in a poised state^8^. Both types of elements, on the other hand, are enriched for acetylation of lysine 27 at histone 3 (H3K27ac) when active^9,10^.

Although the aforementioned features led to the distinction of promoters and enhancers as distinct types, recent work challenged this view, unveiling similarities in their architecture and activity (reviewed in refs^11–13^). Large-scale transcriptome analyses revealed that both promoters and enhancers are bidirectionally transcribed^4,14,15^ and that transcription initiation involves the recruitment of the same transcriptional machinery^16^ (general transcription factors and Pol II). Moreover, although the two types of elements are overall characterized by different chromatin modification profiles, this distinction is not sharp; for example, enrichment of H3K4me3 can also be detected at highly transcribed enhancers^17^. To some extent, the two classes of regulatory elements further show a bivalent functionality, with examples of enhancers acting as alternative promoters^18^ and of promoters enhancing the expression of other genes^19–21^. Despite these observations, which blurred the definition of the two classes of regulatory regions, the association of promoters to long transcripts that are 5’ capped and 3’ polyadenylated still distinguishes these regulatory elements from enhancers, which produce short, generally unstable transcripts^4^.

The extent of stability of nascent transcripts has been linked to the relative enrichment of destabilizing polyadenylation signals (PAS) and stabilizing 5’ splicing (U1) motifs located downstream to the TSS of a transcript. U1 sites, apart from their role in splicing, prevent premature transcript cleavage from cryptic PAS through their binding with the U1 snRNP^22^. Polyadenylation signals proximal to the TSS, on the other hand, have the opposite effect and direct nascent transcripts towards exosome degradation^23^. Unidirectional promoters show a clear enrichment of U1 sites and a depletion of PAS sites in their sense direction relative to their upstream antisense direction, which supposedly limits pervasive genome transcription^24^. The instability of enhancer-associated transcripts is also due to an enrichment of PAS over U1 motifs^17^. This observation led to the prominent hypothesis that changes in the U1-PAS axis might underlie the evolution of new transcripts and, in turn, of new genes^25^. Wu and Sharp proposed that a stepwise gain of U1 sites and loss of PAS motifs in the antisense orientation of previously unidirectional promoters might lead to the emergence of new transcripts that could evolve into new genes upon the acquisition of an open reading frame^25^.

Given the structural and functional similarities between enhancers and promoters, changes in the U1-PAS axis might in principle also lead to a switch in the activity of these regulatory elements. Inheritable mutations at PAS and U1 sites might stabilize enhancer-associated transcripts, thus facilitating their evolution into promoters. Similarly, mutations might destabilize promoter transcription, but not affect the ability of these loci to interact with other regulatory elements and regulate the expression of other genes. In this scenario, one might thus expect to observe orthologous regulatory elements that function as enhancers in one species or organismal lineage but as promoters in another. Interestingly, recent work reported the frequent evolutionary emergence and decay of promoters^26^ and enhancers^27^ in mammals. Although the gain and loss of regulatory elements is largely driven by the insertion and deletion of genomic sequences, such as repetitive elements, many regions align to orthologous loci in other species not showing the same functionality^26,27^, raising the possibility that some of them might have experienced changes in their activity during evolution — a process hereafter referred to as “functional repurposing” (even if the detected events are not necessarily selectively preserved/of phenotypic relevance).

The occurrence of functional repurposing of regulatory elements has been proposed^12,25,28^, and suggestive evidence for the existence of such events has been reported in mammals. We recently described an enrichment of enhancer-associated chromatin marks at mouse loci orthologous to the putative promoters of new rat-specific mRNA-derived gene duplicates^29^ (retrocopies). Moreover, two separate studies reported evidence of 11 mouse long noncoding RNAs (lncRNAs) whose promoter sequences were orthologous to putative human regulatory regions marked by DNase hypersensitivity but not associated to any stable transcript^30,31^. Nonetheless, beyond these potential individual candidates, a thorough investigation of the prevalence of functional repurposing during mammalian evolution and of the underlying molecular mechanisms has been lacking. Here we aim to fill this gap, based on detailed and integrated cross-species analyses of mammalian DNase hypersensitivity, chromatin modification and transcriptional data.

## RESULTS

### Regulatory element repurposing in primates and rodents

As only limited evidence of putative functional repurposing was available from previous studies, we first sought to confirm its occurrence and study its prevalence in mammals. Towards this aim, we defined genome-wide sets of putative enhancers in a mammalian reference species and investigated whether any of these loci were orthologous to putative promoter regions from a closely related species (Fig. 1a) and hence represented candidate repurposed elements – here referred to as “P/E” elements. We focused our work on four species from two mammalian orders: human and rhesus macaque, as representatives of the primate lineage, and mouse and rat from the rodent lineage. We chose these two species pairs for several reasons: first, a large amount of gene expression and chromatin modification data is publicly available for human and mouse, allowing for the annotation of comprehensive sets of regulatory elements from various tissues and developmental stages. Second, the relatively short evolutionary divergence times (25-29 millions of years) between human/mouse and their sister species (i.e., macaque and rat, respectively) facilitates the definition of high confidence orthologous regions for each species pair, thus enabling the comparison of regulatory activities for large numbers of genomic loci. Third, suitable outgroup species (marmoset for the two primates and rabbit for the rodents) with relevant data are available for evolutionary inferences. Finally, both species sets have respective advantages and disadvantages, and therefore the analyses of both datasets allow for overall optimal analyses. For example, while the low mutation rate and resulting high sequence similarity in the primates may allow for higher confidence inferences of early regulatory element evolution, the larger rodent sequence divergence (due to higher mutations rates) and more efficient natural selection during rodent evolution may allow for easier and/or more powerful detection of functional repurposing events.

**Figure 1.**
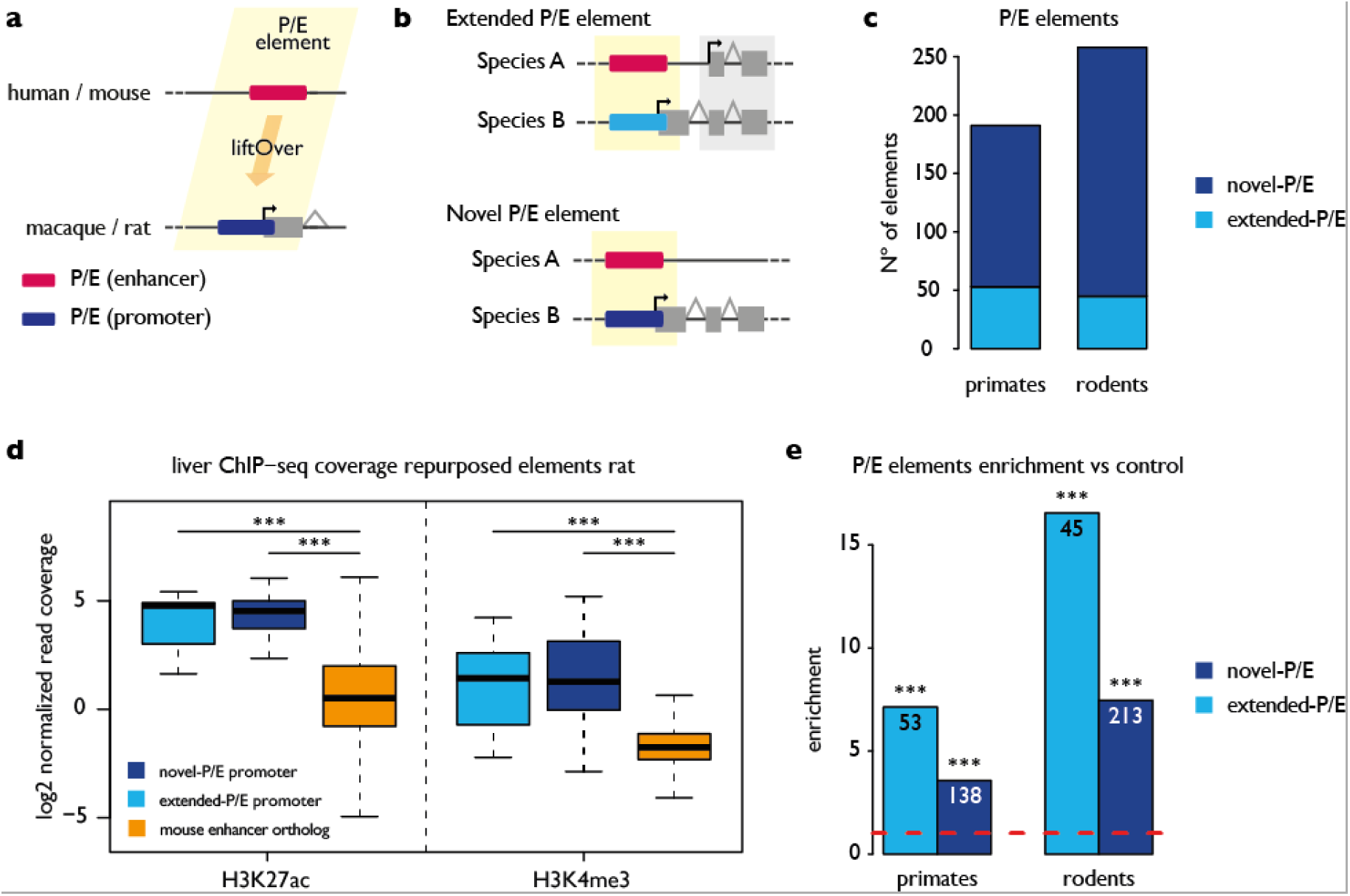
Functional repurposing of regulatory elements in mammals. A) Schematic representation of a P/E element in primates and rodents. B) Types of P/E elements. C) Total number of novel and extended P/E elements detected in primates and rodents. D) H3K27Ac and H3K4me3 ChIP-seq read density from liver (log2 read count normalized by input read count) measured at novel and alternative rat P/E elements compared with rat loci orthologous to mouse enhancers and not associated to any stable TSS. Significant differences (Mann-Whitney *U* test with Benjamini-Hochberg correction): (***) P < 0.001. E) Fold-difference between P/E elements ratio (fraction of human or mouse enhancers corresponding to promoters) and P/random ratio (fraction of human or mouse random unmarked regions corresponding to promoters). The red line indicates no difference between the two ratios. Numbers indicate the number of P/E elements for each group. Significant differences (Chi-squared test with Benjamini-Hochberg correction): (***) *P* < 0.001.

We defined sets of putative promoters as the upstream regions of stably transcribed loci, assembled using both our newly generated as well as recently published^32^ strand-specific RNA-seq data from four adult organs (brain, heart, kidney and liver; see Methods) (Supplementary Data), which yielded between 26,594 and 34,779 promoters for each species (Supplementary Table 1). We then identified putative enhancers in human and mouse (i.e., our reference species for which extensive relevant data are available — see above) by combining transcription, DNase hypersensitivity and histone modification data from the same set of organs. Specifically, we first extracted all DNase hypersensitive sites enriched for H3K4me1 and/or H3K27ac in any of the four organs that were not associated to the TSS of any transcript or enriched for H3K4me3 in any sample from a broad set of tissues and cell types (see Methods). This approach allowed us to annotate a high-confidence set of putative enhancers (Supplementary Table 2) and to avoid the inclusion of potential bivalent elements (e.g. elements characterized by both enhancer and promoter activity in different tissues of the same organism), which would hamper our search for *bona fide* repurposed elements. Additionally, we included in our analysis a second set of putative enhancers, defined using CAGE data from a number of organs and cell lines in human and mouse^4^. Overall, we obtained a total of 117,043 enhancers in human and 131,768 in mouse.

**Table 1:** **Directionality of mammalian repurposing events in liver**

|  | Outgroup species | Converted liver<br>P/E elements | Ancestral liver<br>P/E promoters | Ancestral liver<br>P/E enhancers |
| --- | --- | --- | --- | --- |
| Primates | Marmoset | 53 | 3 (5.6%) | 25 (47.1%) |
| Rodents | Rabbit | 51 | 1 (1.9%) | 16 (31.3%) |

### Widespread functional remodelling facilitated regulatory innovation

After the annotation of putative regulatory regions in our set of species, we assessed the evolutionary conservation of their activity to detect P/E elements. For each human and mouse enhancer, we investigated whether its orthologous locus in the macaque or rat genome, respectively, overlapped the promoter region of a stable transcript (Fig. 1a, Supplementary Table 3). We thus identified 191 P/E elements in primates and 258 elements in rodents (i.e., 449 in total). We subdivided the P/E elements into two distinct categories based on their association to a species-specific (i.e., macaque- or rat-specific) transcribed locus (“novel” P/E) or to a new (species-specific) 5’ exon of a locus transcribed in both species of the respective lineage pair (“extended” P/E) (Fig. 1b-c, Supplementary Table 4). To further confirm the promoter activity of P/E elements in macaque and rat, we inspected their chromatin state using publicly available H3K27ac and H3K4me3 ChIP-seq data from adult liver samples^27^. Consistent with the notion of functional repurposing, P/E elements in both macaque/rat displayed higher H3K4me3 coverage compared to sequences in their genomes that are orthologous to other (non-repurposed) liver enhancers in their sister species (i.e., human/mouse) (Fig. 1d, Supplementary Fig. 1). Our analysis thus revealed the presence of hundreds of P/E elements showing divergent regulatory activities in two major mammalian lineages.

The detection of P/E elements in two closely related species could in principle result from the independent evolution of distinct activities from an ancestral inactive region, rather than from a species-specific repurposing event of an ancestral regulatory region. We therefore investigated whether the presence of enhancer activity at a specific locus would significantly increase the chance of observing promoter activity at the orthologous locus in a closely related species, which would support the repurposing scenario. We defined regions showing no signature of regulatory activity and no overlap with any exonic sequence in human and mouse, and evaluated whether their orthologous regions in the sister species were associated to the TSS of a stable transcript. Only 0.05% (87/187466) of the regions tested in primates and 0.03% (23/78470) in rodents showed this behaviour. These numbers were significantly lower compared to the fraction of P/E elements retrieved in primates (0.19%, >3.56-fold enrichment for novel or extended P/E loci, Chi-squared test, *P* < 10^−15^) and rodents (0.24%, >7.46-fold enrichment, Chi-squared test, *P* < 10^−15^) (Fig. 1e). Although we observed a difference in GC content between the inactive regions and the putative enhancers tested in both species (Supplementary Fig. 2), rat- and macaque-specific promoters were nonetheless more often orthologous to enhancers than to inactive loci with matched sequence composition in their sister species (Supplementary Fig. 3). These data corroborate the hypothesis that P/E elements likely correspond to ancestral regulatory regions that experienced evolutionary changes in their regulatory activity in the last 25-29 millions of years. In other words, our analyses show that ancestral regulatory capacities of genomic sequences facilitated regulatory innovation and suggest that the “de novo” origination of regulatory activity is rare.

**Figure 2.**
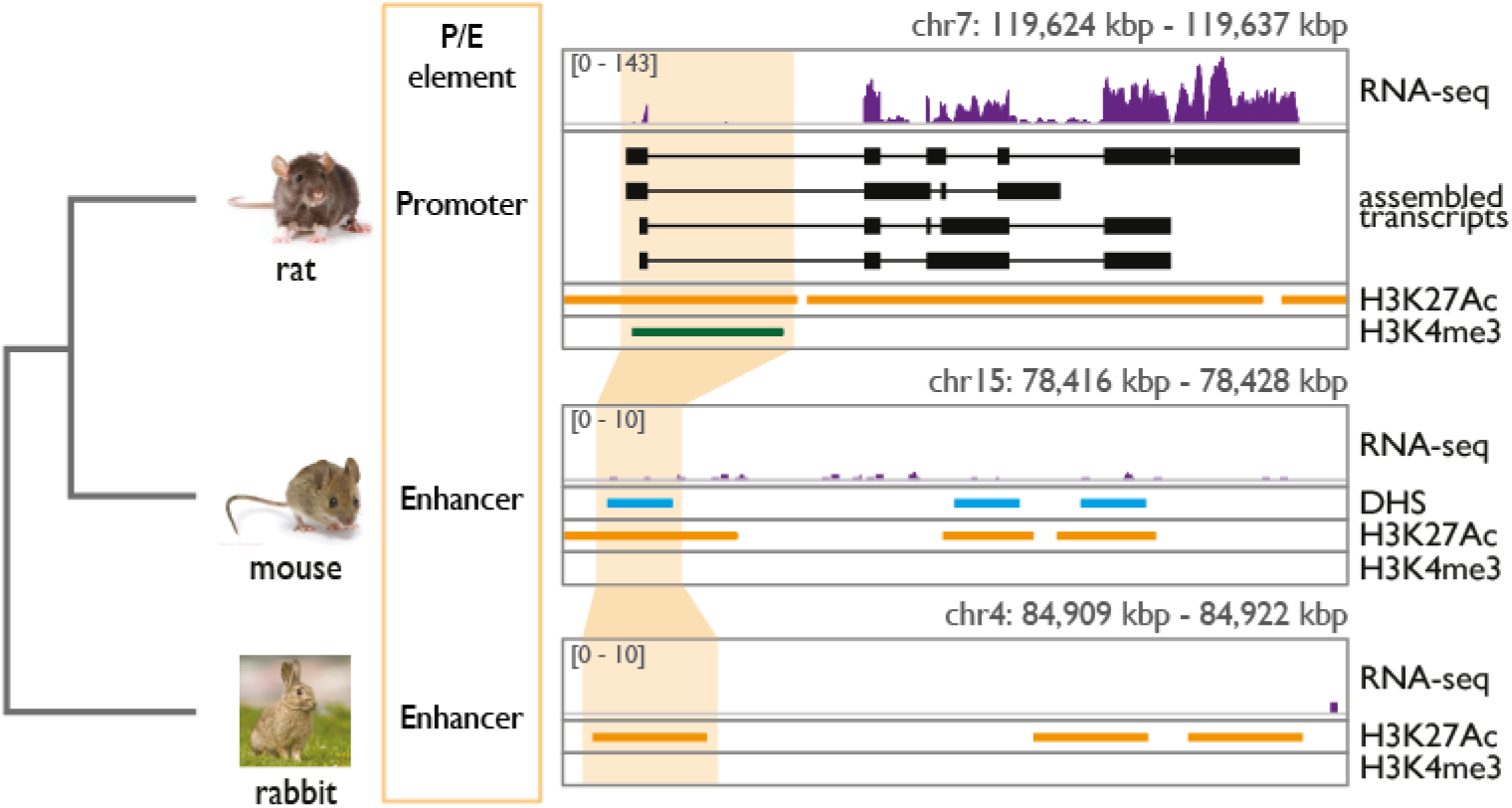
Repurposing of an ancestral glires enhancer. The coordinates of a rodent P/E element, with enhancers activity in mouse and promoter activity in rat, are projected onto the rabbit genome (orange bars). The syntenic regions in rabbit overlaps an H3K27ac peak but no H3K4me3 peak. The presence of a transcript in rat and absence of transcripts in mouse and rabbit are evident based on the RNA-seq tracks. For each species (from top to bottom) are shown: the RNA-seq coverage from liver; the assembled transcripts (only in rat); the liver DHS peaks (only in mouse); the liver H3K27ac and the H3K4me3 peaks from Villar et al. (2015).

**Figure 3.**
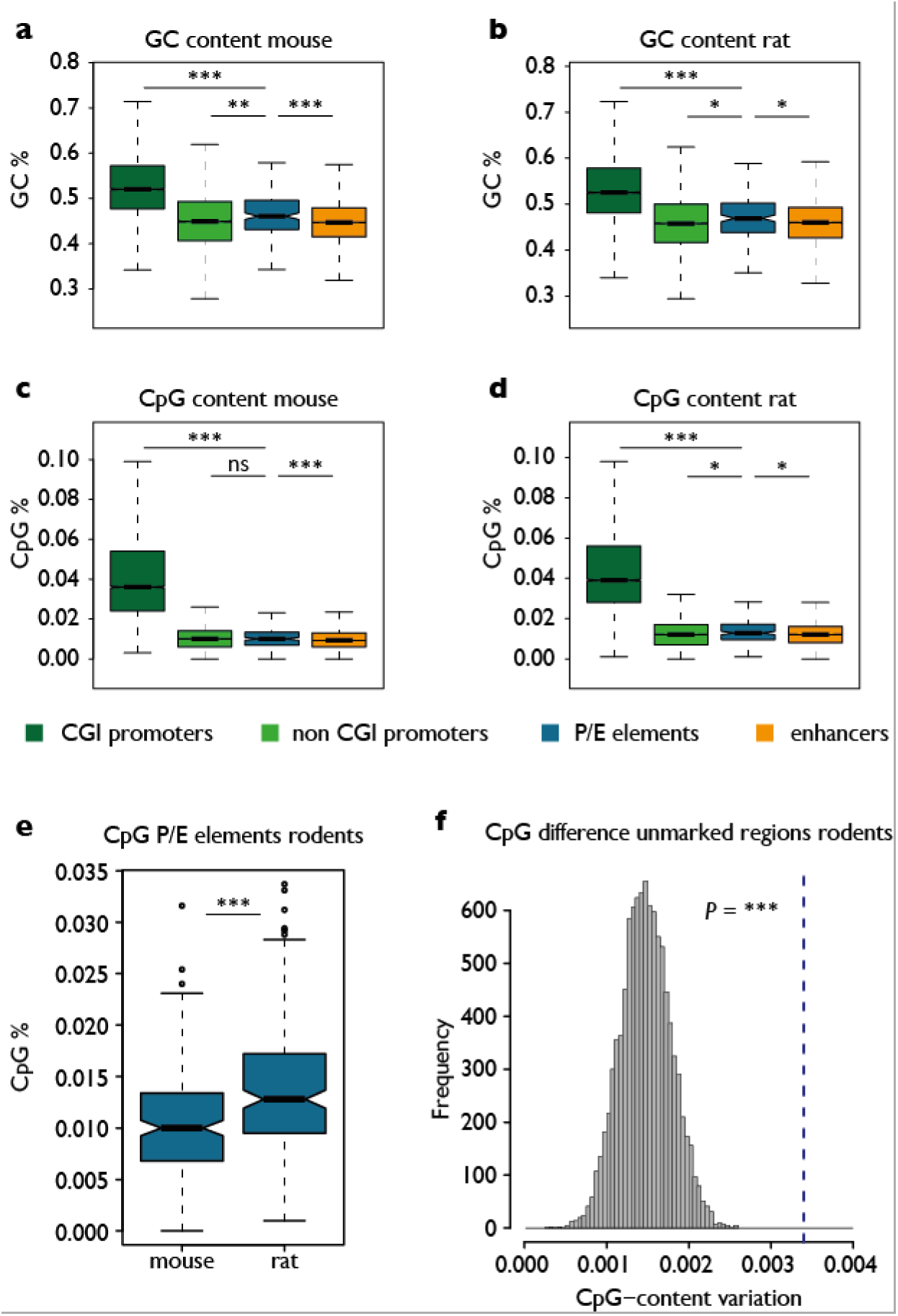
Nucleotide composition of P/E elements. A-D) Distribution of GC and CpG content for different classes of regulatory elements in mouse and rat. Significant differences (Mann-Whitney *U* test with Benjamini-Hochberg correction): (***) P < 0.001; (**) P < 0.01; (*) P < 0.05; (n.s.) P ≥ 0.05. E) Distribution of CpG dinucleotide frequency in orthologous rodent P/E elements. F) A number of orthologous rodent unmarked regions equal to the rodent P/E elements in panel D) was resampled 10,000 times. The histogram shows the distribution of mean CpG density difference between the resampled orthologous unmarked regions. The dashed line indicates the mean CpG density difference between rodent orthologous P/E elements. Significant differences (resampling test): (***) P < 0.001.

### Ancestral enhancers are the main source of functional repurposing

The retrieval of hundreds of lineage-specific P/E elements allowed us to investigate at a broad scale the directionality of regulatory activity changes; that is, to define whether an ancestral enhancer evolved into a promoter, or vice versa. We thus investigated the presence of regulatory activity associated to regions orthologous to P/E elements in an outgroup species, in order to infer their ancestral state. Using ChIP-seq and transcription data to annotate putative regulatory elements in adult marmoset liver, we observed that 47% (25 of 53) primate P/E elements with activity in liver and aligned to the marmoset genome correspond to orthologous putative enhancers in this outgroup species, whereas only ≈5% overlapped a promoter (Fig. 2, Table 1). Similarly, ≈31% of rodent P/E elements correspond to putative enhancers in rabbit, while only ≈2% overlapped a promoter (Table 1). The much higher fraction of ancestral P/E elements with enhancer activity strongly suggests that most repurposed elements correspond to ancestral enhancers that recently evolved species-specific promoter activities.

To evaluate whether the bias in the directionality of the repurposing events could be explained by the higher evolutionary turnover of enhancers compared to promoters^27^, we compared the rates of repurposing and loss of activity for ancestral enhancers and promoters in both lineages (Methods, Supplementary Fig. 4). In the primate lineage, we identified 2,256 ancestral enhancers and 1,370 ancestral promoters that lost their activity in macaque. In contrast, 25 ancestral enhancers and 3 ancestral promoters changed their activity in human or macaque (≈5-fold enrichment, Fisher’s exact test, *P* < 10^−2^). These observations suggest that the higher rate of enhancer repurposing cannot be simply explained by the higher turnover rate (i.e., reduced selective preservation) of these elements compared to promoters. The lack of a statistically significant similar pattern in rodents (≈5-fold enrichment, Fisher’s exact test, *P* = 0.08; Supplementary Fig. 4) is probably explained by genomic/evolutionary differences between the two sets of species. That is, the considerably larger divergence time of the rodent species and outgroup (mouse/rat–rabbit divergence time: ≈80 million years, my) compared to that in primates (human/macaque-marmoset: ≈42 my), the higher mutation rates in glires (the clade including rodents and lagomorphs), and/or the larger long-term effective population sizes (i.e., more efficient natural selection) in glires may obscure actual rates of activity turnover in this lineage (e.g., elements that lost regulatory activity may have been completely lost or have diverged too much to be aligned across species; mainly beneficial events have been retained; there may have been multiple turnovers at the same locus).

**Figure 4.**
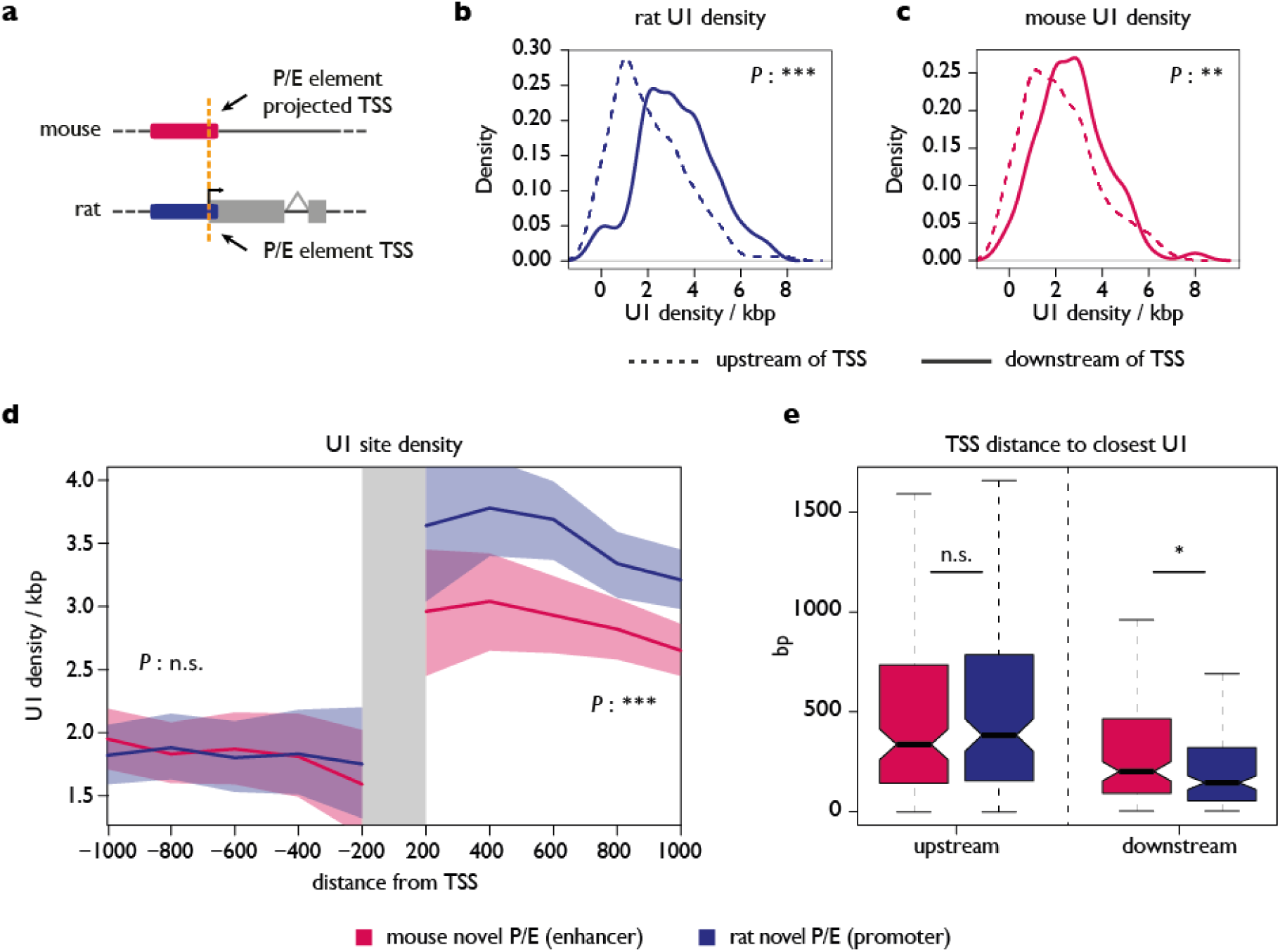
U1 site distribution at P/E elements. A) Schematic representation of an orthologous P/E element. The orange line depicts the position of the TSS in rat and of the projected location in mouse. BC) Distribution of U1 site density per kb upstream (dashed line) and downstream (continuous line) of the TSS of transcripts associated to novel P/E elements in rat (B) and mouse (C). Significant differences (Mann-Whitney *U* test with Benjamini-Hochberg correction): (***) P < 0.001; (**) P < 0.01. D) Cumulative density of U1 sites up- and downstream of novel P/E-associated TSSs in rodents. Lines represent the mean U1 density (per kb) over 200, 400, 600, 800 and 1000 nt-long windows from the TSS, shaded areas represent 95% confidence intervals. Significant differences (Mann-Whitney *U* test with Benjamini-Hochberg correction): (***) P < 0.001; (n.s.) P ≥ 0.05. E) Distribution of up- and downstream distances of the closest U1 site from each novel P/E-associated TSS in rodents. Significant differences (Mann-Whitney *U* test with Benjamini-Hochberg correction): (*) P < 0.05; (n.s.) P ≥ 0.05.

Our analysis also revealed that between 25 and 34 (47% and 66% in primates and rodents, respectively) of the P/E elements active in liver did not bear any signature of activity in the same organ from the outgroup species, and we reasoned that some P/E elements might be active in a different organ. We thus used RNA-seq data to define the promoters of transcribed regions in marmoset and rabbit, and then determined the fraction of primate and rodent P/E elements corresponding to a promoter in any of the adult organs investigated. This analysis showed that only 11 of the 178 marmoset regions orthologous to a primate P/E element (6.1%) overlapped the TSS of a transcript and therefore likely corresponded to an ancestral promoter. An even lower fraction (4/129, 3.1%) of rodent P/E elements corresponded to promoters in rabbit. These results further confirmed that only a limited number of repurposing events involved ancestral promoter elements, supporting the hypothesis that evolutionary changes in regulatory activity in these two mammalian lineages disproportionately involved the evolution of new promoters from ancestral enhancer elements.

### P/E elements have distinct sequence compositions

Large-scale surveys have demonstrated that the sequence composition of mammalian enhancers closely resembles that of promoter regions that do not overlap with CpG islands (CGIs), but differences between distinct types of enhancers have been reported^4^. For example, mammalian enhancers with broad spatial expression patterns are characterized by a higher overlap with CGIs compared to other enhancers. Due to the peculiar change in activity of P/E elements, we therefore investigated whether their sequence composition would be distinct relative to other regulatory regions. P/E elements in both human and mouse (i.e., the species with P/E enhancer activity) have an overall lower GC and CpG content when compared with CGI-associated promoters, in agreement with the general lack of association between CpG islands and enhancers (Fig. 3a-b, Supplementary Fig. 5a-b). Surprisingly, we observed an overall higher GC content in P/E elements compared to both non CGI-associated promoters and other enhancers across all species (except for macaque enhancers), indicating that the sequence composition distinguishes P/E elements from other regulatory sequences – regardless of their activity (Fig. 3a-b, Supplementary Fig. 5ab). A similar trend was observed for the CpG frequency, with significantly higher content of this dinucleotide in P/E elements compared to enhancers and to non-CGI promoters across species (except for non-CGI promoters in human and mouse) (Fig. 3c-d, Supplementary Fig. 5c-d), reinforcing the distinction of this class of regulatory elements from other active loci.

### Sequence compositional changes probably contributed to repurposing

Next we sought to trace whether compositional changes may underlie functional shifts of P/E elements. Notably, the CpG content of regulatory elements has been proposed to influence their transcriptional output^33,34^. CpG islands are usually associated to the promoter of broadly and highly expressed genes^1^, where they favour gene expression by creating a nucleosome-free environment^33,34^. We then asked whether P/E elements with promoter activity were associated to higher GC- and CpG-contents compared to their orthologous regions. To do so, we compared the difference in GC and CpG frequencies between orthologous P/E elements with that observed between orthologous inactive regions (likely not subjected to selective sequence constraint) to control for potential global differences in the sequence composition of each species pair. We found that although rat P/E elements showed somewhat higher GC-content compared to their orthologous sequences in mouse, the observed difference was not significantly stronger than that of the control regions (Supplementary Fig. 6), in agreement with the previously reported higher genome-wide GC-content in rat^35^. By contrast, we noted that the total content of CpG dinucleotides in rat P/E elements increased significantly compared to the control regions (Fig. 3e-f), indicating that the activity turnover of P/E loci is mirrored by a change of CpG frequency in this lineage. In primates, the GC-content did not differ significantly between the orthologous P/E elements, whereas the frequency of the CpG dinucleotides was significantly higher in macaque (i.e., for P/E elements with promoter activity) compared to human (i.e., for P/E enhancers) (Supplementary Fig. 7). Notably, the small enrichment in both GC and CpG content measured in macaque was statistically significant when compared to the control regions (Supplementary Fig. 7). This result reflects the slightly lower GC content of the macaque genome compared to the human one^35^. Overall, the reported effect of CpG content on transcription^33,34^ (see above) together with the higher CpG-content in the species where the P/E element acts as a promoter suggest that specific changes in nucleotide composition contributed to the regulatory repurposing of P/E elements both in rodents and in primates.

### Distinct U1 motif and PAS signal distributions at P/E elements

While promoters and enhancers both have the inherent capacity to promote transcription, only promoters generate stable transcripts^17^. In mammals, this difference between the two types of regulators seems to be mainly directed by specific signals downstream of the TSS. It has been shown that U1 sites are commonly enriched downstream of promoters and depleted in the antisense orientation as well as around enhancers, whereas PAS generally follow the opposite trend^4,24^. Owing to their potential role in transcription, we compared the distribution of U1 signals and PAS around orthologous novel P/E elements (Fig. 1b, lower panel). For each element, we extracted U1 and PAS motifs up- and downstream of the TSS of their associated transcript in macaque and rat, as well as for the corresponding orthologous regions in the respective sister species (Fig. 4a). In both macaque and rat, as expected, given the promoter function of P/E elements (and association with stable transcripts) in these species, we observed a higher density of U1 sites downstream of the TSS compared to the antisense orientation and a weak but significant opposite trend for the PAS motifs (Fig. 4b, Supplementary Fig. 8). In human and mouse, where annotated P/E elements have enhancer properties (and no stable transcription is detectable), we found no difference in PAS distribution around each P/E element but noted a significantly higher density of U1 sites downstream of the projected TSS (Fig. 4c, Supplementary Fig. 8). Therefore, P/E elements not associated to stable transcripts are nonetheless characterized by a U1/PAS environment with mixed features compared to typical promoters and enhancers, which — together with their unique sequence composition (see above) — may predispose them to repurposing during evolution.

### Evolutionary changes in the U1/PAS axis

Evolutionary changes in the U1/PAS axis have been proposed as a mechanism underlying the emergence of new transcribed loci that may be selectively preserved and thus form new genes^25^. However, so far, evidence supporting this hypothesis has been lacking. We therefore took advantage of our dataset of orthologous P/E element pairs to test whether evolutionary changes in the U1 and PAS motif distribution around these loci might underlie their regulatory activity transformation. By comparing the distribution of U1 sites surrounding orthologous P/E elements in rodents, we found that their density increased significantly over 1 kilobase (kb) downstream of the TSS of promoter-associated rat transcripts with respect to the orthologous non-transcribed regions in mouse enhancers (mean of 3.21 vs. 2.65 U1 sites per kb, Wilcoxon’s test, Benjamini-Hochberg corrected *P* < 10^−4^, Fig. 4d), whereas no significant difference was observed in the corresponding upstream regions. When comparing the PAS distribution for the same regions, we observed only a weak decrease in PAS density downstream of the rat TSSs (mean of 1.33 vs 1.64 PAS sites per kb, Wilcoxon’s test, Benjamini-Hochberg corrected *P* < 10^−2^, Supplementary Fig. 9) and no difference in the antisense orientation. We further observed a significantly shorter distance separating the TSS from the closest downstream U1 site in rat promoters compared to the orthologous mouse enhancers (Wilcoxon’s test, Benjamini-Hochberg corrected *P* < 10^−2^, Fig. 4e). Finally, U1 sites preceded a PAS downstream of a P/E-associated TSS in rat in 84.5% of the cases, compared to 71.6% for the orthologous mouse enhancers (Chi-squared test, *P* < 10^−2^). On the other hand, we found no significant difference in U1 and PAS motif distributions around P/E elements in primates (Supplementary Fig. 10), probably due to the low sequence divergence and resulting lack of power^36^. Overall, our data revealed evolutionary shifts in the distribution of U1 sites and, to a lesser extent, PAS motifs, which mirrored the presence or absence of stable transcripts (i.e., promoter or enhancer activity) at P/E loci in rodents. Thus, changes in the U1/PAS axis indeed seem to contribute to the origination of promoters (from enhancers) and, as a consequence, the emergence of new transcribed loci in mammals.

## DISCUSSION

Mammalian promoters and enhancers share many similarities in their chromatin architecture, and — apart from a minor fraction of bivalent elements^20^ — these regulatory loci are best distinguished based on the stability of their associated transcripts^17^. This suggests that small changes in the DNA sequences underlying or surrounding regulatory regions could redefine their activity. In our work, we provide strong support for this hypothesis by identifying hundreds of mammalian elements that experienced an evolutionary turnover in their regulatory activity, and by tracing specific sequence changes that accompanied this process.

Previous attempts to identify evolutionarily repurposed regulatory sequences uncovered 11 mouse lncRNA promoters orthologous to putative enhancer elements in human^30,31^. The low number of candidate elements identified likely resulted from the long evolutionary distance separating the two species. Moreover, the divergent activity in these cases might not necessarily result from a repurposing event, but could rather have evolved independently in the two lineages. In our study, we shed light on the repurposing process by focusing on the comparison of more closely related species (within the primate and rodent lineages, respectively) and by using inactive genomic regions as controls to confirm that the divergent activity of P/E elements likely results from an actual evolutionary switch in their function. The hundreds of P/E elements uncovered here indicate that the evolutionary turnover of regulatory elements activity is much more extensive than could have been estimated based on the human/mouse comparisons. Moreover, the number of elements uncovered in our work for the investigated species is likely to represent an underestimate. For example, we conservatively excluded from our analysis a large number of putative bivalent regulatory elements (characterized by enhancer and promoter activity in the same species) in order to maximize the confidence in detecting true turnover events, which may have led to the removal of many real enhancers characterized by H3K4me3 enrichment^17^, and we had to focus on only one of the two species (i.e., the reference species human and mouse, where substantial chromatin data are available) for the initial enhancer annotation (i.e., a bidirectional analysis would likely have uncovered around twice the number of events detected here). In any event, our work indicates that the repurposing of regulatory elements activity is a widespread and previously unappreciated process shaping the mammalian regulatory landscape.

The investigation of P/E element activity in outgroup species revealed that most turnover events seem to involve the repurposing of ancestral enhancer elements into species-specific promoters, an observation that we show not to be solely explained by the higher evolutionary turnover of enhancers (due to reduced purifying selection) compared to promoters in mammals^27^. It should be noted that almost half of the alignable P/E elements in each lineage had no detectable activity in the outgroup species. This is likely due to the relatively large evolutionary distance that separates our core set of species from their evolutionary outgroups, indicating that the regulatory activity of these loci might either have emerged after the split of the outgroup lineages or that it was lost during the evolution of the outgroup species lineages. Although we cannot exclude that the inferred directionality of the repurposing process is influenced by the lack of definition of the ancestral state for part of the P/E loci, such a scenario is unlikely to fully explain the biased enhancer-to-promoter conversion pattern, because the higher rate of enhancer turnover reduces the likelihood of detecting enhancers conserved in more distantly related species, which should in principle disfavour the detection of enhancer-to-promoter turnover events. Our results therefore suggest the existence of differences in repurposing potential between enhancers and promoters, which could involve their underlying DNA sequence and/or aspects of their chromatin composition. Future work, involving more closely related species or different populations of the same species, will be necessary to further explore the biased directionality of the repurposing process and uncover its mechanistic bases.

The sequence analysis of P/E elements revealed sequence features that distinguish these loci from other regulatory loci, and it provided initial evidence for the potential mechanisms behind the repurposing process. Notably, the high GC and CpG content could make P/E loci particularly prone to drive the expression of neighbouring sequences, for example through the recruitment of CpG binding proteins such as Cfp1^37^. This protein is known to deposit H3K4me3 marks over the bound sequence^38^, which in turn seems to favour transcription through different mechanisms^39,40^. Moreover, recent work showed how CpG sites favour promoter over enhancer activity in a massively parallel regulatory element assay in mouse^41^. The significantly higher CpG content of P/E elements with promoter activity strongly suggests that the fixation of nucleotide substitutions contributed to the turnover events by increasing (or decreasing) the density of this dinucleotide, leading to the creation or disruption of specific motifs that altered transcriptional capacities.

Moreover, U1 site density shifts also seem to be involved in the repurposing process. A higher number and a higher proximity of U1 sites characterize the region downstream of the TSS of P/E-associated transcripts, compared to their transcriptionally inactive orthologous regions. U1 sites are thought to promote transcript stability in mammals, suggesting that changes in the distribution of these motifs might be responsible for the stabilization or destabilization of P/E associated RNAs. On the other hand, it is unclear whether the distribution of polyadenylation signals (PASs) had a significant influence on the turnover process. Although PASs are slightly depleted downstream of the TSS compared to the upstream region specifically in macaque and rat (i.e., the species used to assess P/E promoter activity), we found no significant differences in PAS distribution between orthologous P/E elements, suggesting that, at least in this context, variation in U1 site distribution could be sufficient drive to the repurposing process.

Finally, from an evolutionary perspective, this work sheds novel light on the process of the birth and death of new genes/transcripts. Our results provide solid evidence for the widespread emergence of new stable transcripts from enhancer elements that may, potentially, represent new genes. Our work therefore provides initial strong support to the prominent hypothesis by Wu and Sharp^25^ and highlights functional repurposing as an notable mechanism underlying molecular innovation in mammals. Future work that focuses on the functional consequences of turnover events will unveil the overall impact of the repurposing process on mammalian phenotypic evolution.

## MATERIALS AND METHODS

### RNA-seq data production and processing

We generated 78 single-end strand-specific RNA-seq libraries from brain, heart, kidney and liver samples for 6 mammals (human, macaque, marmoset, mouse, rat and rabbit; Supplementary Table 5) using the Illumina TruSeq Stranded mRNA LT kit according to manufacturer instructions. Libraries were sequenced using the Illumina HiSeq 2000 to produce 100 nucleotide (nt) reads. The resulting transcriptome data were combined with our recently published transcriptome data^32^.

RNA-seq reads were aligned to the assembled genomes (Ensembl release 73) of their corresponding species using TopHat2^42^ (version 2.1.0) using the following parameters: -a 8 -i 40 -I 1000000 --read-realign-edit-dist 0 --microexon-search. Genome indexes needed for the alignment were generated using Bowtie2^43^ (version 2.2.4). Aligned reads from all replicates of each organ (totalling on average >100 million mapped reads) were used to reconstruct transcripts through a genome-guided de novo transcriptome assembly using StringTie^44^ (version 1.2.0) with the following parameters: -j 5 –g 50. Assembled transcripts from each organ were then merged using Cuffmerge^45^ to define a unique set of transcripts. Expression levels (measured in FPKM) of the assembled transcripts were calculated with Cuffnorm^45^ (version 2.2.1); we considered as stable all transcripts with a mean FPKM > 1 across replicates from the same organ and length > 1000 nt.

### ChIP-seq and DNase-seq data processing

The chromatin data used in our studies derive from different sources. DNase, H3K4me3, H3K4me1 and H3K27ac data for mouse brain, heart, kidney and liver (core dataset) were obtained from the Mouse ENCODE database^46^ (Supplementary Table 6). DNase, H3K4me3, H3K4me1 and H3K27ac data from human brain, heart, kidney and liver were obtained from the ENCODE database^47^ or from the human Epigenome Roadmap database^48^ (Supplementary Table 6). For both species, we also downloaded H3K4me3 data from additional adult and developmental samples from the same databases (extended dataset) (Supplementary Table 6). All processed data corresponding to an older genome assembly version from mouse (mm9) were converted to the newest version (mm10) using LiftOver^49^. As data were processed in different ways, we applied a common approach to have comparable datasets. Specifically, we downloaded processed peaks (in narrowPeak format) from multiple replicates of all samples, and subsampled the top 20’000 (for H3K4me3 data) or top 80’000 peaks (for H3K4me1 and H3K27ac), ranked based on their peak score; all DNase hypersensitive site (DHS) peaks from each sample were retained as their numbers did not differ significantly across samples. We created organ/tissue specific sets of H3K4me3, H3K4me1 and H3K27ac peaks by considering loci shared by at least three replicates from each organ (or by both replicates if only two samples were available), except when peaks were already derived from merged samples, as for the adult mouse organs. We finally resized the peaks to 1,000 nt centred on the summit of the peak (or on the middle of the peak when the summit was not available).

### Definition of regulatory and random inactive regions

In each species, we defined as promoters the 1,000 nt located upstream of a stable transcript. Putative enhancers in human and mouse were initially defined as DHSs overlapping an H3K27ac and/or an H3K4me1 peak. The resulting set of enhancers was further filtered to exclude loci located closer than 1,000 nt from any H3K4me3 peak from any organs/tissues (including the extended dataset), or overlapping the 1000 nt upstream region or the exons of any (stable or unstable) transcript. We further downloaded enhancer sets defined using CAGE from human and mouse^4^ (“permissive enhancers phase 1 and 2” from http://fantom.gsc.riken.jp/5/datafiles/latest/extra/Enhancers/). These loci were subjected to the same filtering process described above, and then included in the final list of putative enhancers.

To define the set of random inactive regions, we sampled from the human and mouse genome 1.5 million non-overlapping 1000 nt loci, and then removed from this list all loci mapping closer than 1000 nt from: a) any DHS or any H3K4me1, H3K4me3 or H3K27ac peak from all organs from the core and extended dataset; b) any exon from an (stable or unstable) or annotated (from GENCODE)^50,51^ transcript; c) any high identity (95%) segmental duplication or repeat element annotated in the UCSC database^52^.

### Definition of orthologous regulatory and random inactive loci and detection of turnover events

Coordinates of all regulatory and random regions from human and mouse were converted on the macaque and rat genome, respectively, using LiftOver^49^ (with -minMatch=0.6) to define their orthologous loci. A two-way liftOver conversion (species A -> species B -> species A) was adopted to avoid errors in the orthology definition that may result from genomic duplication events. We defined as P/E elements all human and mouse enhancers whose orthologous loci in macaque and rat, respectively, overlapped the 500 nt upstream of the TSS of a stable transcript. As a control, we identified all random inactive regions in human and mouse whose orthologous sites in their sister species overlapped the 500 nt upstream of the TSS of a stable transcript, and then compared the frequency of these loci with the frequency of P/E elements in each core sample with a Chi-squared test. P/E elements were analysed separately whether the associated transcript in macaque or rat corresponded to a new isoform of a transcribed locus present in the sister species (extended P/E element) or to a newly transcribed locus (novel P/E element). Shared transcribed loci between orthologous species were defined by the overlap of the orthologous sequences of macaque/rat transcripts (determined using liftOver) with reconstructed or GENCODE annotated transcripts in human/mouse.

After analysing the sequence composition of the enhancers and random regions (with an ortholog in the sister species) used in the aforementioned analysis, we observed a significant difference in GC-content between the two sets. To control for the GC-content effect, we resampled sets of enhancers and random regions with a similar GC distribution. To this aim, we considered only enhancers with a GC-content lower than 46% (for human) or 41% (for mouse). These thresholds, roughly corresponding to the mode of the GC distribution of the enhancers set in the two species, were chosen as they represented the maximum value below which the GC distribution of random regions overlapped completely the GC distribution of the enhancers. Then, we further subsampled up to 30,000 enhancers from each species (in order to have similar numbers of regions tested in both lineages), and for each locus we selected a random region with the most similar GC content. With this approach, we obtained sets of enhancers and random regions with statistically similar GC distributions and used these data to compare the frequency of P/E enhancers to that of random regions orthologous to promoters.

### ChIP-seq analysis of P/E elements

To further support the promoter functionality of P/E promoters in rat and macaque, we compared their H3K4me3 and H3K27ac profiles to those of other regulatory elements using liver ChIP-seq data obtained from Villar et al.^27^. In both species, the enhancer set corresponded to the putative enhancers projected from their sister species, whereas promoters were defined as previously described (see “Definition of regulatory and random inactive regions”). Finally, we compared the average H3K4me3 and H3K27ac coverage, normalized using the average input coverage, between all classes of regulatory elements with a Mann-Whitney *U* test.

### Polarization of turnover events

To define the directionality of the turnover events, we evaluated the presence of putative promoters or enhancers in regions syntenic to P/E elements active in liver in marmoset and rabbit. In marmoset and rabbit, enhancers corresponded to H3K27ac peaks not overlapping any H3K4me3 peak or the 1000 nt upstream and the exons of any (stable or unstable) assembled transcript; promoters were defined as the described before. Macaque and rat liver P/E element coordinates were converted in their outgroup species genome using LiftOver (with -minMatch=0.4), and we then evaluated the overlap between the converted coordinates and the annotated regulatory regions.

To estimate the rate of regulatory element loss, we identified the orthologous loci of liver promoters and enhancers from human and mouse in the corresponding outgroup species (marmoset and rabbit, respectively), and kept only those loci that showed the same activity in the two species. Specifically, ancestral promoters had to be associated (e.g. overlap the 500 nt upstream of the TSS) to a stable liver transcript; ancestral enhancers had to overlap an H3K27ac peak and not be associated to a promoter or an H3K4me3 peak. These loci were then aligned on the macaque/rat genome, and the loss of activity in these species was defined by the lack of stable transcription or H3K27ac peaks for ancestral promoters or enhancers, respectively.

To estimate the fraction of ancestral promoters corresponding to P/E elements, we further calculated the overlap of primate and rodent P/E elements coordinates converted in marmoset and rabbit, respectively, with the putative promoter of any stable transcript (from any organ).

### Sequence composition of regulatory elements

We extracted and compared the GC content [using the nuc tool from BEDTools^53^] and CpG dinucleotides frequency of different classes of regulatory elements in human and mouse using a Mann-Whitney *U* test. The same features were compared between orthologous P/E elements in both lineages with a Wilcoxon signed rank test. Finally, we estimated whether the magnitude of the evolutionary change in GC content or CpG frequency at P/E regions was higher than that measured at random regions (defined above) to control for global skews in sequence composition. This was done by resampling 10000 times from the set of random regions the same number of P/E elements, and then comparing the mean GC and CpG difference in P/E elements with the distribution of differences from the resampled set.

### U1/PAS motif composition of regulatory elements

We determined the genome-wide location of U1 and PAS sites with the scanMotifGenomeWide tool from HOMER^54^ (version 4.7). To compare the distribution of the U1 and PAS motifs around P/E elements, we considered the leftmost TSS of all P/E associated transcripts in macaque and rat and projected their location in the corresponding sister species using BLAT^55^. We considered only novel P/E elements, given that U1/PAS motifs from downstream transcripts in extended P/E loci might have conflated the U1/PAS signal in mouse and human. We compared the density of U1 and PAS motifs over 1 kb up- and downstream of the P/E-associated TSS in each species using a Wilcoxon signed-rank test. The same approach was used to compare the density of these motifs between sister species. The proximity of the closest U1 or PAS site up- or downstream of the P/E-associated TSSs was calculated using closestBed from BEDTools^53^.

### Software used

All processing of genomic coordinates was performed using tools from BEDTools suite^53^ (version 2.19.1). All statistical analysis were performed using R^56^. In-house code used to perform all analyses is available upon request.

## DATA ACCESS

Raw and processed data sets from this study have been submitted to the NCBI Gene Expression Omnibus (GEO; http://www.ncbi.nlm.nih.gov/geo/) (*accession numbers pending*).

## ACKNOWLEDGEMENTS

We thank K. Harshman and the Lausanne Genomics Technology Facility for high-throughput sequencing support; I. Xenarios and the Vital-IT computational facility for computational support; the Kaessmann group and Ana Claudia Marques for helpful discussions. The overall research project was supported by a grant from the European Research Council (Grant: 615253, OntoTransEvol) to H.K. F.N.C. was supported by an SNSF Early Postdoc.Mobility fellowship (P2LAP3_171808).

## CONTRIBUTIONS

F.N.C. and H.K. designed the study. F.N.C. conducted the analysis. A.L. and J.H. performed the experiments. F.N.C, M.W. and H.K. wrote the manuscript.

